# Halophytic endophytes provide transferable immune functions for crop resilience

**DOI:** 10.64898/2026.07.21.739774

**Authors:** Vassiliki A. Michalopoulou, Stefanos K. Soultatos, Savvas Paragkamian, Nikolaos P. Arapitsas, Theoni Margaritopoulou, Maria-Frantzeska Triviza, Sofia Kosenaki, Eleni Kalogeropoulou, Emilia Markellou, Emmanouil A. Markakis, Panagiotis F. Sarris

## Abstract

Halophytic microbiomes represent an untapped reservoir of stress-adapted microbial functions, yet whether these functions can be transferred across ecological boundaries to enhance crop immune resilience remains unknown. Here we show that *Kushneria*, a halophyte-associated genus with broad environmental resilience, confers robust disease protection in tomato and cucumber plants against fungal and oomycete pathogens despite lacking direct antagonistic activity. Comparative genomics across 26 genomes revealed extensive biosynthetic novelty and conserved catabolic clusters associated with rhizosphere competence, providing a genomic framework for persistence of Kushneria in saline environments. Multi-omics profiling associated protection with a distinct host immune regulatory response, characterized by strong induction of core NLR receptors, including a ZAR1 paralogue. This response was further linked to a previously uncharacterized long noncoding RNA, Solyc10r048010.1, connected to the induced NLR network. Our findings establish halophytic endophytes as field-validated and transferable modulators of crop immunity and identify a candidate regulatory RNA–NLR axis associated with *Kushneria*-induced disease resistance.

## Introduction

Climate-driven salinization is emerging as a major constraint on agricultural productivity, compromising both crop performance and the efficacy of conventional microbial inoculants^1–4^. Halophytic plants host microbial communities adapted to extreme osmotic stress, yet the roles of these endophytes in crop systems remain insufficiently defined. Microbiomes adapted to extreme environments may provide an untapped reservoir of microbial functions that have evolved to maintain host performance under stress and may be transferable across ecological contexts. Their biological relevance may depend not only on natural host association, but also on the transferability and efficacy of these stress-adapted functions under agricultural conditions.

Members of the genus *Kushneria* (*Halomonadaceae*) are frequently isolated from saline environments and halophytic hosts^5–9^ and have been associated with plant growth promotion under hypersaline conditions^10,11^. Whether halophyte-associated endophytes can transfer ecologically evolved functions into glycophytic crops, and whether such interactions reveal conserved principles of microbiome-mediated immunity, remains poorly understood. Microorganisms originating from saline ecosystems possess physiological and metabolic traits that may support activity under conditions that destabilize conventional crop-associated microbes, but whether such traits confer advantages for disease management or immune modulation has not been established.

Beneficial bacteria suppress disease either through direct antagonism or by activating host immunity^1^^2^. Although immune priming by beneficial microbes is well documented^12,13^, it remains unknown whether extreme-environment microbiomes constitute a reservoir of immune-associated functions that can be transferred across ecological boundaries into crop hosts. Understanding how such transferable host-protective functions are perceived and integrated by crop immune systems also remains largely unresolved. In particular, the coordinated roles of intracellular immune receptors, such as NLRs, and post-transcriptional regulators, including non-coding RNAs, represent a critical knowledge gap. Furthermore, it remains unclear whether halophyte-derived endophytes trigger distinct immune responses in crop hosts. This question is particularly relevant under the global effect of climate change and the phenomenon of water salinization, where many established inoculants lose efficacy while halophytic endophytes may retain functional stability^14,15^.

Here, we characterized halophyte-derived *Kushneria* endophytes using an integrated genome-to-field framework combining comparative genomics, multi-omics and field validation. We show that *Kushneria* isolates confer broad-spectrum disease protection in tomato and cucumber despite lacking direct antagonistic activity, demonstrating that halophyte-associated endophytes can transfer immune-associated functions across ecological boundaries into glycophytic crop systems. Comparative genomics revealed extensive biosynthetic novelty together with conserved rhizosphere competence traits consistent with persistence under saline conditions, while multi-omics profiling identified a distinct immune reprogramming response marked by induction of core NLR receptors and a candidate long non-coding RNA–NLR regulatory framework. Together, these findings provide proof of concept that immune functions associated with halophytic microbiomes remain functional following transfer into glycophytic crops.

## Results

### Isolation and genomic characterization of endophytic *Kushneria* spp

Endophytic bacteria were isolated from four halophytic plant species on the island of Crete and initially identified by 16S rRNA gene sequencing. Nine *Kushneria* isolates were selected for whole-genome sequencing and functional characterization. Using a hybrid long- and short-read strategy, we assembled complete circular genomes for all isolates (Ext. Data Table 1). Genome sizes were ∼3.6 Mb with G+C contents of 58–59%, consistent with previously described *Kushneria* species^5–7^, although isolate SRL383 exhibited a slightly higher value. Several isolates also carried additional replicons, including putative plasmids, and SRL376 harbored a third replicon (Ext. Data Table 2).

Taxonomic assignment using FastANI^16^ revealed varying degrees of relatedness to available reference genomes. SRL376 and SRL377 were most closely related to *Kushneria phyllosphaerae* (GCF_900312995.1)^7^, sharing 95.07% identity, whereas SRL383 showed 98.0% identity to *Kushneria indalinina* (GCF_003385845.1)^5,17^. The remaining isolates were most similar to strain 4-2069 (GCF_947497025.1), with ANI values of 91.57–93.33% (Ext. Data Table 1). As these values fall below commonly used species delineation thresholds, several isolates likely represent previously undescribed species within the genus *Kushneria*.

### Phylogenetic and pangenome analysis of *Kushneria* isolates

To resolve the taxonomic relationships among the isolates, we performed whole-genome comparisons using digital DNA–DNA hybridization (dDDH). Clustering of 26 available *Kushneria* genomes yielded 17 species-level groups, with the nine SRL isolates distributed across five species and five subspecies clusters (Fig. 1, Ext. Data Table 3). Among these, only SRL383 was assigned to *Kushneria indalinina*, whereas the remaining isolates likely represent previously undescribed species or subspecies within the genus (Ext. Data Table 4).

**Figure 1.**
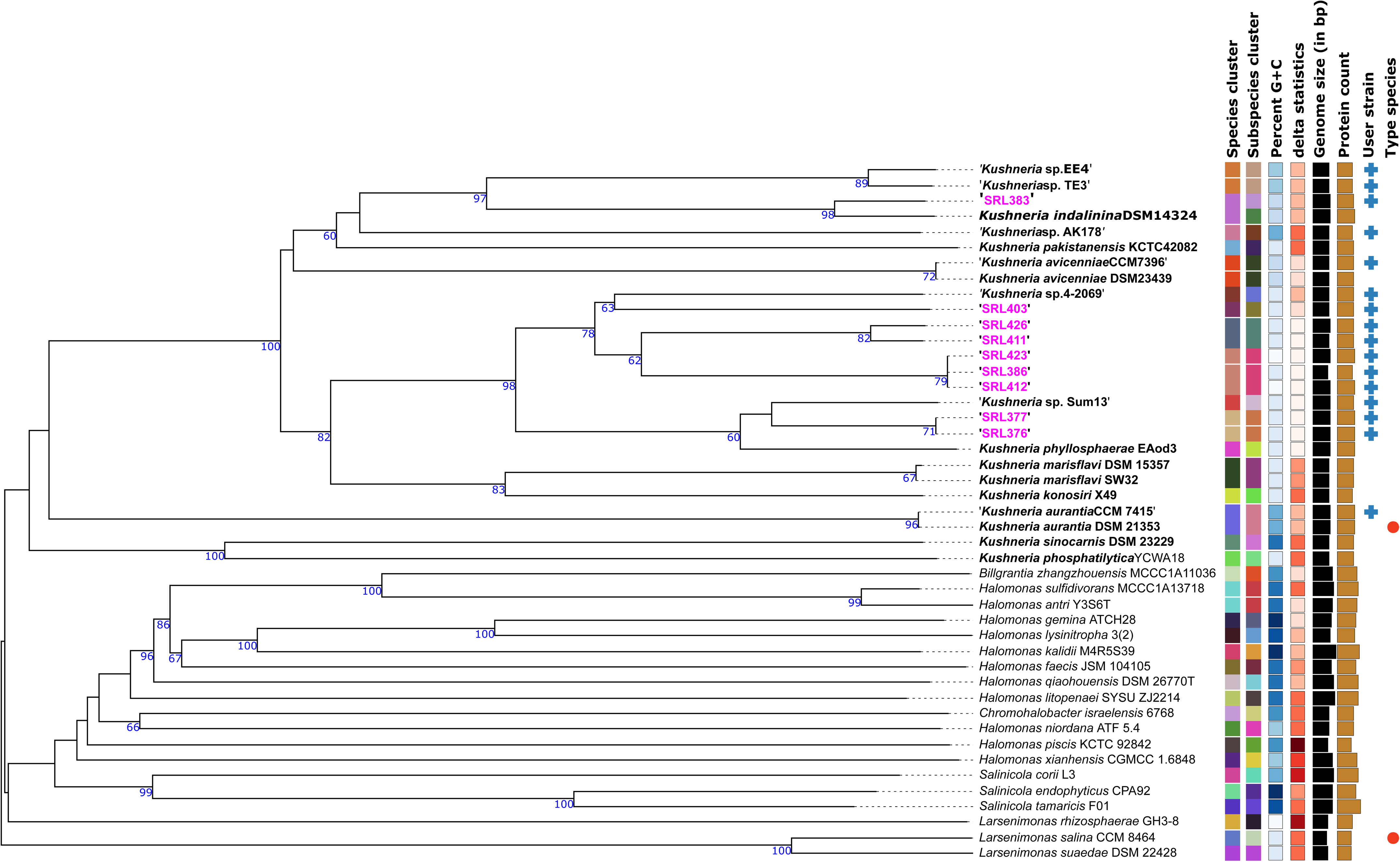
A phylogenetic tree of all *Kushneria* spp. genomes. Phylogenetic tree showing the placement of the nine endophytic *Kushneria* assemblies (purple labels) among publicly available type-strain genomes (45 genomes total). The tree was inferred with FastME 2.1.6.1^47^ from GBDP distances calculated from whole-genome sequences. Branch lengths are scaled according to GBDP distance formula d5. Numbers above branches indicate GBDP pseudo-bootstrap support values >60% from 100 replications (average branch support: 73.8%). The tree was midpoint-rooted^45^.

To place the isolates within the broader diversity of the genus, we constructed a genus-wide pangenome using all 26 available genomes (as of 05/01/2026). The pangenome exhibited substantial diversity, with only 17.1% of gene clusters (GCs) constituting the core genome (Fig. 2, Ext. Data Table 5). Phylogenomic analysis separated the genus into two major lineages: one comprising *K. sinocarnis*, *K. phosphatilytica* and *K. aurantia*, and a second containing four subclades that included all SRL isolates. Within this latter lineage, SRL376 and SRL377 clustered with *K. phyllosphaerae*; SRL383 clustered with *K. indalinina* and *K. pakistanensis*; and SRL403, SRL411 and SRL426 clustered with *K. marisflavi* and *K. konosiri*. In contrast, SRL386, SRL412 and SRL423 formed a distinct clade (Fig. 2).

**Figure 2.**
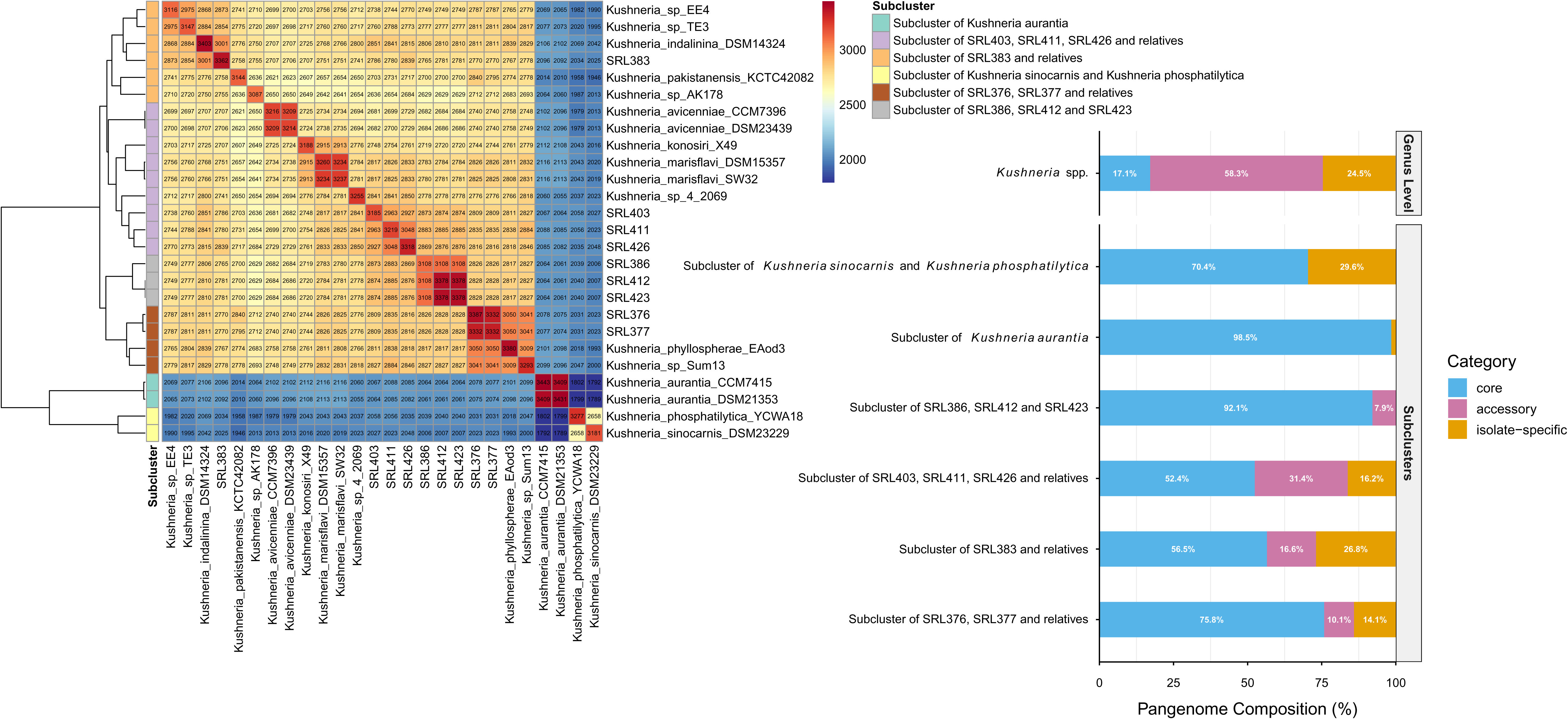
Pangenome analysis of selected isolates with their closest relatives. Heatmap and bar plot showing the pangenome of *Kushneria* spp. (26 genomes), including the nine endophytic isolates sequenced in this study. Core, accessory and isolate-specific gene clusters (GCs) are shown at the genus level and for each subclade. The number of GCs per genome is indicated within each cell, and the color gradient represents the proportion of shared GCs between genomes. A subcluster annotation column is shown on the left for easier interpretation of pangenome structure, corresponding to the clustering on the x- and y-axes (see Extended Data Fig. 1 for additional information on shared GCs among subclusters).

Comparisons of core, accessory and isolate-specific GCs further resolved relationships among closely related isolates (Ext. Data Fig. 1A–D). SRL383 shared ∼80% of its GCs with *K. indalinina*, but FastANI and dDDH analyses support its classification as a distinct subspecies (Ext. Data Table 6, Ext. Data Fig. 1C). SRL412 and SRL423 shared an identical core GC complement, consistent with their assignment to the same strain and their common origin from the leaves of *Euphorbia paralias*, yet differed from SRL386 isolated from *Tetraena alba* leaves by 270 isolate-specific GCs (Ext. Data Fig. 1A). Similarly, SRL376 and SRL377 shared 98.4% of their GCs, supporting their classification within the same species (Ext. Data Table 6). Both these isolates were obtained from leaves of *Atriplex halimus*. Although SRL403, SRL411 and SRL426 initially clustered together, SRL403 shared only 78.7% of its core GCs with SRL411 and SRL426, indicating that it represents a distinct species-level lineage within this group (Ext. Data Table 6). Notably, SRL403 was obtained from the same sample as SRL386; *Tetraena alba* leaves.

### Primary and secondary metabolic potential of *Kushneria* spp

Analysis of metabolic module completeness across the *Kushneria* pangenome revealed patterns broadly consistent with phylogenetic relationships. SRL383 grouped more closely with *K. pakistanensis* than with *K. indalinina*, further supporting its classification as a distinct subspecies (Ext. Data Table 7, Ext. Data Fig. 2A).

Genome mining identified a conserved repertoire of nine biosynthetic gene clusters (BGCs), including arylpolyene, ectoine, redox-cofactor, RiPP-like, hserlactone, NRPS, NRPS-independent siderophore (NI-siderophore) and two terpene clusters, across complete *Kushneria* genomes (Fig. 3A, Ext. Data Table 5). Although these BGCs were broadly conserved, NI-siderophore loci varied among subclades, with different lineages encoding distinct siderophore types (Ext. Data Table 8). Comparison with the MiBIG database showed that most predicted BGCs lacked close matches to characterized clusters, indicating substantial unexplored biosynthetic diversity within the genus. Exceptions included ectoine and selected NI-siderophore clusters, which showed moderate similarity to clusters from *Methylophaga thalassica* and *Pantoea agglomerans*, respectively (Ext. Data Fig. 2B, Ext. Data Table 8).

**Figure 3.**
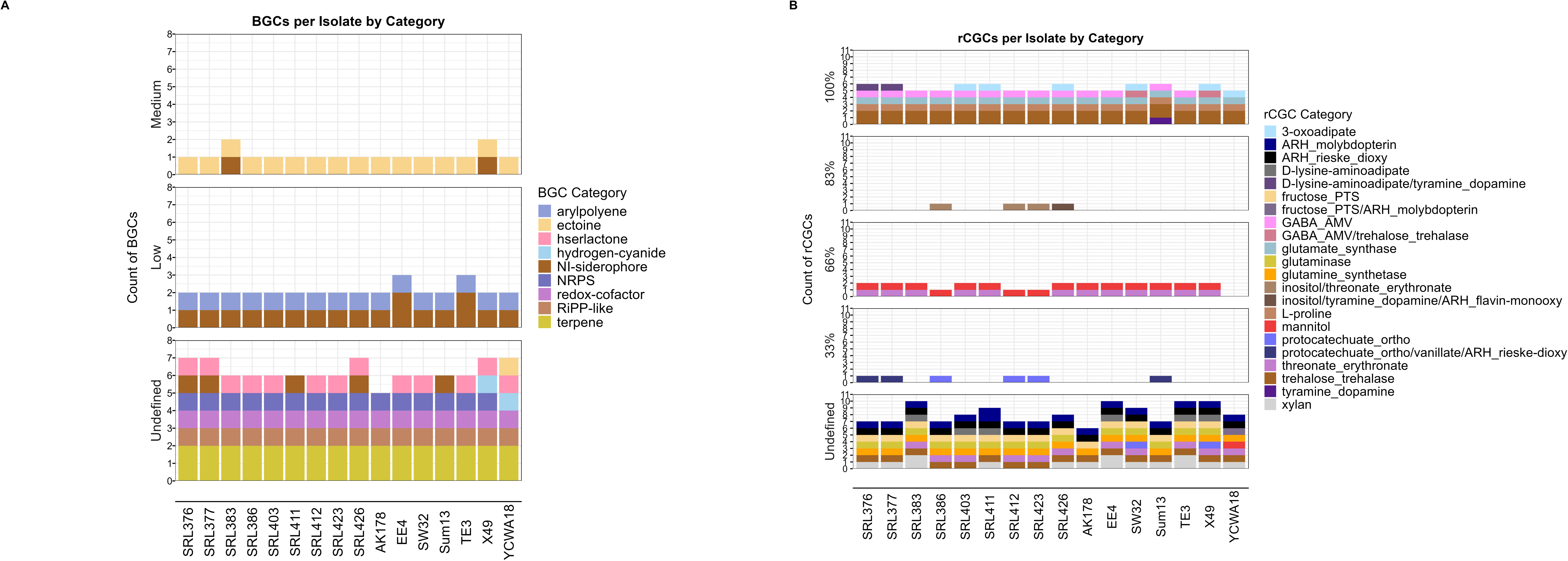
Distribution of the BGCs and the rCGCs across different *Kushneria* spp. **A.** Stacked bar plots showing the number of biosynthetic gene clusters (BGCs) identified per isolate, grouped by similarity confidence to known BGCs (Medium, Low, Undefined) as assigned by antiSMASH. Only complete genomes were analyzed, including the nine isolates sequenced in this study. BGCs were further grouped into nine categories: arylpolyene, ectoine, hserlactone, hydrogen-cyanide, NI-siderophores, NRPS, redox-cofactor, RiPP-like and terpenes. “Undefined similarity” refers to clusters lacking close matches to known BGCs in the antiSMASH KnownClusterBlast database. See Extended Data Table 8 for sequence similarity comparisons and closest MIBiG matches, and Extended Data Fig. 2B for total BGC counts per isolate. **B.** Stacked bar plots showing rhizosphere catabolic gene clusters (rCGCs) grouped by similarity confidence (100% to undefined), as assigned by rhizoSMASH. rCGCs were categorized into 16 single-function groups: 3-oxoadipate, ARH-molybdopterin, ARH-Rieske-dioxygenase, D-lysine–aminoadipate, fructose-PTS, GABA-AMV, glutamate synthase, glutaminase, glutamine synthetase, L-proline, mannitol, protocatechuate-ortho, threonate/erythronate, trehalose-trehalase, tyramine/dopamine and xylan. Additional dual or triple-function rCGCs included D-lysine–aminoadipate/tyramine–dopamine, fructose-PTS/ARH-molybdopterin, GABA-AMV/trehalose-trehalase, inositol/tyramine–dopamine/ARH-flavin-monooxygenase and protocatechuate-ortho/vanillate/ARH-Rieske-dioxygenase. See Extended Data Fig. 2C for total rCGC counts per isolate.

To evaluate traits potentially associated with plant colonization, we analyzed complete genomes using rhizoSMASH^18^. Twelve core rhizosphere-associated catabolic gene clusters (rCGCs) involved in the utilization of root-derived compounds were identified across the genus (Fig. 3B). Conserved functions included pathways for L-proline, mannitol, glutamine, glutamate and trehalose metabolism, whereas variation was observed in xylan and threonate/erythronate utilization pathways. Most genomes additionally encoded clusters associated with GABA metabolism, fructose uptake, glutaminase activity and ARH-molybdopterin-dependent functions (Fig. 3B). These conserved osmoadaptive profiles provide a genomic rationale for the structural stability and persistence of halophytic isolates within glycophytic rhizospheres.

Despite broad conservation of rhizosphere-associated functions, differences in genomic organization were observed among closely related strains. For example, SRL376 and SRL377 encoded a protocatechuate/vanillate/ARH-Rieske-dioxygenase region comprising both single and hybrid catabolic clusters, whereas the corresponding loci in the related strain Sum13 were organized as a single hybrid cluster (Ext. Data Table 9).

### Broad-spectrum disease suppression by *Kushneria* spp. in crop hosts

To assess the disease-protective potential of *Kushneria* isolates, we evaluated their effects against multiple fungal and oomycete pathogens. *In vitro* assays revealed no antagonistic activity against *Fusarium oxysporum* f. sp. *radicis-lycopersici* (*Forl*), *Botrytis cinerea* and *Phytophthora nicotianae*, except in the case of SRL376 but with no statistical significant reduction of radial growth of *B. cinerea* (Ext. Data Fig. 3).

Despite the absence of direct inhibition, several isolates significantly reduced disease development *in planta*. In tomato, SRL412 and SRL426 suppressed disease progress, resulting in a reduced area under the disease progress curve (AUDPC) following *Forl* infection (Fig. 4A). SRL386 reduced *B. cinerea* disease progression in both whole-plant and detached-tissue assays (Fig. 4B, C), whereas SRL376 decreased the crown rot lesion size caused by *P. nicotianae* by ∼50% (Fig. 4D), although it did not reduce the AUDPC following *B. cinerea* infection (Ext. Data Fig. 4). Conversely, SRL426 increased *B. cinerea* disease (Ext. Data Fig. 4). Disease suppression was not restricted to a single host species: in cucumber, inoculation with SRL376, SRL412 and SRL426 isolates significantly reduced powdery mildew (*Podosphaera xanthii*) severity in both whole-plant and leaf-disc assays (Ext. Data Fig. 5).

**Figure 4.**
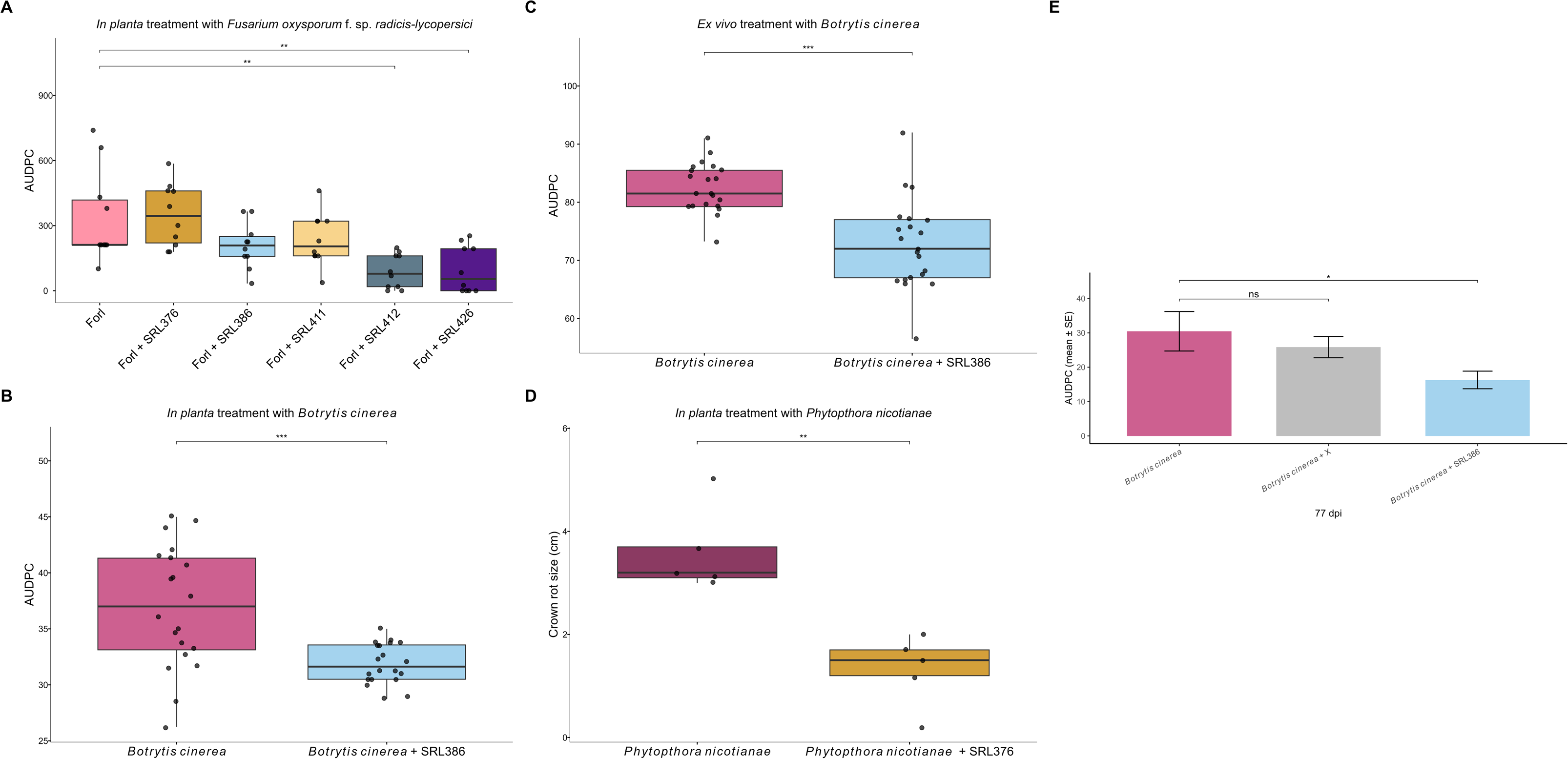
*In planta* and/or *ex vivo* bioassays against different phytopathogens in *Solanum lycopersicum*. Disease-protection assays of endophytic SRL isolates in *S. lycopersicum*. **A.** Boxplot showing the AUDPC caused by *Fusarium oxysporum* f. sp. *radicis-lycopersici* (Forl) infection of tomato plants treated with SRL376, SRL386, SRL411, SRL412 or SRL426 (pathogen alone as control). AUDPC at 21 dpi showed significant reductions for Forl + SRL412 and Forl + SRL426 (n = 10, p < 0.01). **B, C.** Boxplots showing the AUDPC caused by *Botrytis cinerea* infection of tomato plants (B) and fruits (C) treated with SRL386 (pathogen alone as control). AUDPC was significantly lower for Bc + SRL386 in both assays (plants: n = 20, p < 0.001; fruits: n = 21, p < 0.001). See Extended Data Fig. 4C for additional isolates tested. **D.** Boxplot showing the crown rot size caused by *Phytophthora nicotianae* infection of tomato plants treated with SRL376 (pathogen alone as control). Crown-rot size was significantly reduced in Phn + SRL376 (n = 5 per treatment, p < 0.01). **E.** Barplot showing the AUDPC caused by natural *B. cinerea* infection in field-grown tomato plants treated with SRL386 or a commercial *Bacillus amyloliquefaciens* product (X). Only SRL386 significantly reduced AUDPC at 77 days post treatments (n = 100, p < 0.05).

To determine whether these effects extend to agricultural settings, we conducted a field trial evaluating the efficacy of SRL386 against *B. cinerea*, including a commercial *Bacillus amyloliquefaciens* product (X) as a reference. Across a 77-day evaluation period, SRL386 significantly reduced disease development (AUDPC), whereas the commercial product did not produce a statistically significant reduction under the tested conditions (Fig. 4E).

Together, these results demonstrate that selected *Kushneria* isolates provide protection against diverse fungal and oomycete pathogens across multiple crop hosts despite exhibiting no direct antagonistic activity *in vitro*. The broad-spectrum nature of this protection, together with its reproducibility under field conditions, supports the hypothesis that *Kushneria* confers transferable host-mediated immune functions across crop systems.

### Host immune reprogramming associated with Kushneria-mediated protection

Given the absence of direct antagonistic activity *in vitro*, we investigated whether disease suppression by *Kushneria* is associated with host immune activation. To maximize mechanistic resolution within a single, robust host–microbe model, we focused our multi-omics profiling on isolate SRL386, selected due to its consistent protective effects under both controlled and field conditions. As a control for “priming-like” responses, we compared its effects with those of *Streptomyces* sp. SRL307, which did not reduce *B. cinerea* pathogenicity in tomato (Ext. Data Fig. 4).

#### Defense-associated transcriptional responses

SRL386 treatment significantly altered the expression of defense-associated marker genes, including *PR-1*, *LoxD* and *PDF1A*. PR-1 and LoxD remained elevated between 24 and 72 h post-inoculation, whereas PDF1A showed a transient increase at 24 h (Ext. Data Fig. 6A). In contrast, SRL307 induced limited changes, including a transient reduction in PR-1 and increased LoxD expression at 24 h (Ext. Data Fig. 6B). For SRL386 treatment, NPR1 and WRKY70 were upregulated from 24 to 72 h, AOS increased at 48 h, MYC2 at 72 h, and WRKY33 remained elevated after one week (Ext. Data Fig. 6A). These results are consistent with a broader and more sustained transcriptional response than the SRL307 strain.

#### Transcriptome-wide reprogramming associated with SRL386 treatment

RNA sequencing revealed extensive transcriptomic reprogramming in tomato leaves following SRL386 treatment (48 hpi), with 1,567 differentially expressed genes (DEGs; |log₂FC| ≥ 1), comprising 716 upregulated and 851 downregulated genes (Ext. Data Fig. 7A, Ext. Data Table 10). In contrast, SRL307 induced comparatively few transcriptional changes (Ext. Data Fig. 7B, Ext. Data Table 10).

KEGG enrichment analysis showed that SRL386-upregulated genes were enriched for plant hormone signaling pathways and caffeine metabolism, whereas downregulated genes were associated with ribosome-related functions and photosynthesis (Fig. 5A, B). Opposite trends were observed following SRL307 treatment (Ext. Data Fig. 7C, D). Gene set enrichment analysis (GSEA) further identified enrichment of lipid metabolic pathways, including α-linolenic acid and linoleic acid metabolism, fatty-acid degradation and autophagy-related processes (Fig. 5C).

**Figure 5.**
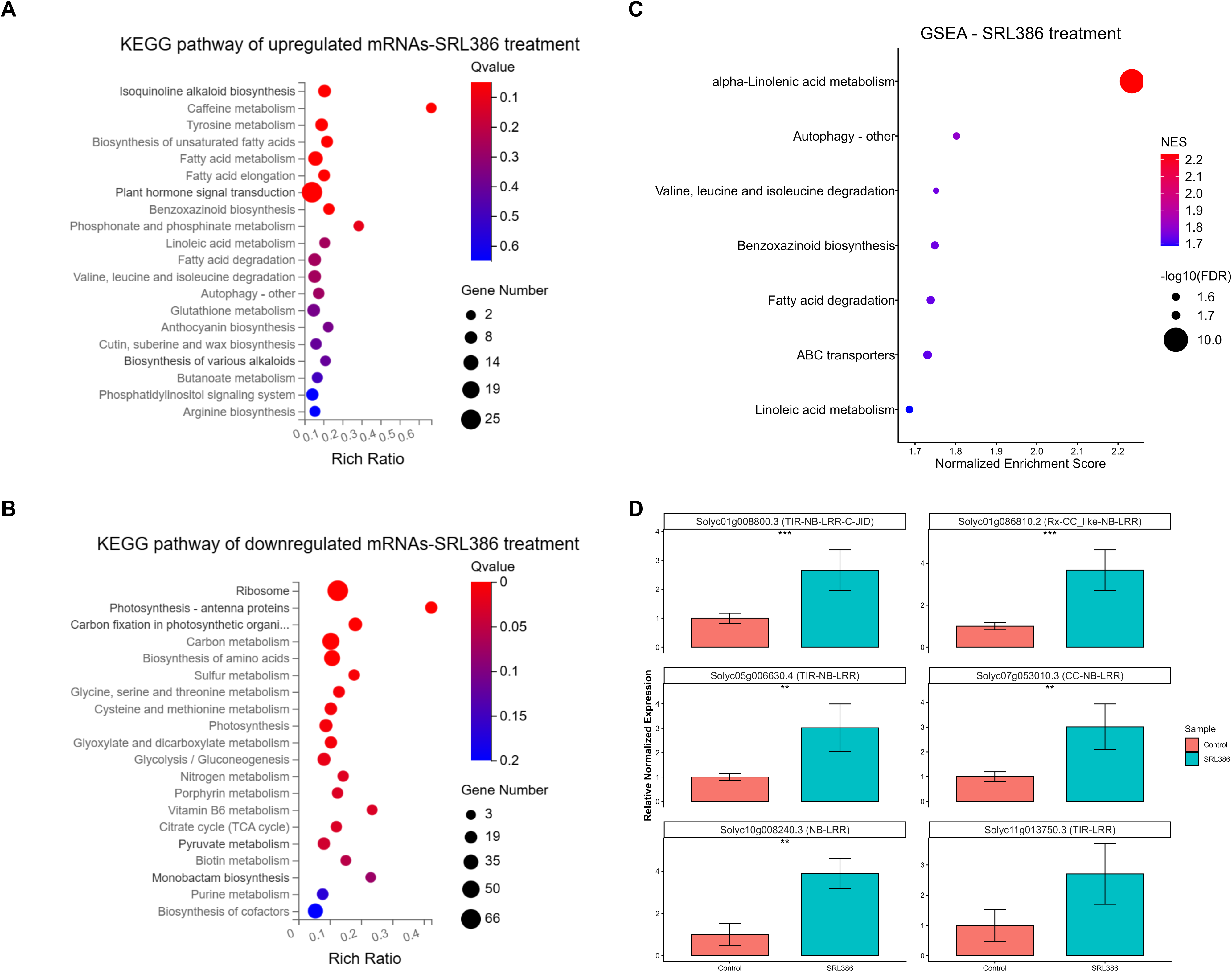
Transcriptomic profile of *Solanum lycopersicum* plants treated with SRL386 KEGG pathway enrichment analysis of upregulated. (**A**) and downregulated (**B**) genes following SRL386 treatment. Pathways with Q ≤ 0.05 are shown. Bubble size represents the number of genes per pathway; color indicates enrichment significance. Upregulated genes were enriched for plant hormone signaling and caffeine metabolism, whereas downregulated genes were enriched for ribosome-related functions and photosynthesis. **C.** Gene set enrichment analysis (GSEA) showing KEGG pathways with FDR < 0.05. α-Linolenic acid metabolism was the top significantly enriched pathway. **D.** qRT-PCR validation of seven NLR genes upregulated at 48 h post-treatment. Two are TIR-NLRs; Solyc01g008800.3 contains a C-JID integrated domain (SMART^78^). Two are CC-NLRs, including the ZAR1-like Solyc07g053010.3. Solyc10g008240.3 contains NB and LRR domains, and Solyc11g013750.3 contains TIR and LRR domains. Solyc08g005510.3 showed very low transcript abundance and was detected in only one replicate. Significance: *p < 0.05, **p < 0.01, ***p < 0.001.

Notably, seven nucleotide-binding leucine-rich repeat (NLR) genes were significantly upregulated following SRL386 treatment and validated by qRT-PCR (Fig. 5D). This response occurred despite the absence of detectable Type III secretion systems (T3SSs) or Type III effectors(T3Es) in the sequenced *Kushneria* genomes (Ext. Data Table 11).

#### Regulatory RNA networks associated with host responses

To investigate regulatory components underlying these transcriptional changes, we profiled microRNAs (miRNAs) and long non-coding RNAs (lncRNAs). This analysis identified 47 and 21 differentially expressed (DE) miRNAs and lncRNAs, respectively (Ext. Data Fig. 8A, B, Ext. Data Table 10). We next predicted their respective targets. Specifically, for lncRNAs, we identified targets among both *cis*- and *trans*-acting genes, which were subsequently integrated with the DE mRNA dataset to construct a regulatory network comprising 673 nodes and 832 edges. (Fig. 6).

**Figure 6.**
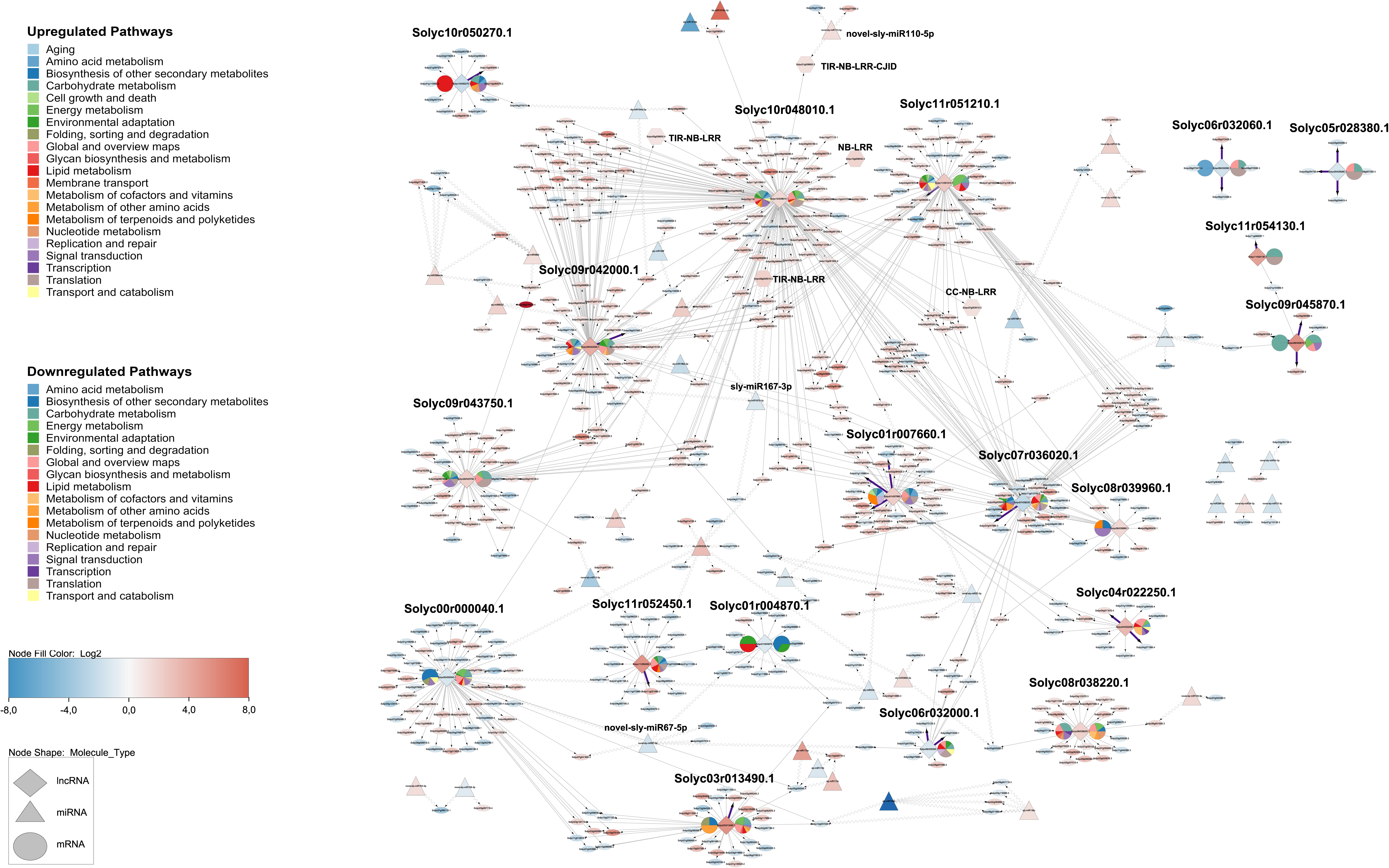
Regulatory *in silico* network of transcriptomic changes by the treatment of tomato plants with *Kushneria* sp. SRL386 Network analysis of differentially expressed (DE) lncRNAs (diamonds) and their DE cis- and trans-mRNA targets (ellipses), and DE miRNAs (triangles) and their DE targets in SRL386-treated *S. lycopersicum* plants. Edges represent predicted interactions: lncRNA–cis-mRNA (purple, thicker lines), lncRNA–trans-mRNA (light blue), miRNA–lncRNA (sinewave) and miRNA–mRNA (sinewave). Node color indicates log₂ fold change. Each lncRNA contains one or two pie charts showing KEGG pathway terms of up-regulated (left) and/or down-regulated (right) mRNA targets (NA pathways omitted). Hexagons denote upregulated NLRs predicted to be all targeted by a common lncRNA, the Solyc10r048010.1. Network was constructed in Cytoscape v3.10.4.

Among the DE lncRNAs, Solyc00r000040.1, a chloroplast-associated lncRNA targeted by 3 distinct miRNAs, was strongly downregulated and was predicted to target genes exclusively *in trans*. Most of its upregulated targets were involved in biosynthesis of other secondary metabolites. Reversely, most of the downregulated targets were associated with energy metabolism and global and overview maps (Fig. 6, Ext. Data Table 12).

The lncRNA Solyc10r048010.1 displayed the highest network connectivity and was predicted to interact with 154 trans-target genes, including five of the seven induced NLRs, among them the ZAR1 paralogue *SlZAR1-sis-1*. Additionally, both isoforms of this lncRNA were predicted to be targeted by sly-miR167b-3p, represented in the network as double edges (Fig. 6).

Most mRNAs were predicted to be targeted by one single miRNA, although several exhibited multiple targeting relationships involving genes associated with aging, transcription, translation, protein folding, degradation and transport (Ext. Data Fig. 8C, Ext. Data Table 12).

Collectively, these results indicate that SRL386 treatment is associated with extensive host transcriptional reprogramming involving defense-related pathways, NLR induction and coordinated changes in regulatory RNA networks.

## Discussion

Our findings support a framework in which extreme-environment microbiomes function as transferable reservoirs of crop-relevant immune functions. Rather than restricting microbial discovery to crop-associated communities, exploring stress-adapted ecosystems could reveal resilient functional traits that remain effective under agricultural and climate stress.

Halophytic endophytes represent an underexplored reservoir of stress-adapted microbial functions with potential relevance for crop resilience under increasing environmental stress, yet their capacity to modulate plant immunity has remained largely undefined. Here, we show that *Kushneria* isolates derived from halophytic hosts confer broad-spectrum disease protection in tomato and cucumber despite lacking direct antagonistic activity. To our knowledge, this represents the first demonstration that members of the genus *Kushneria* function as immune modulators in glycophytic crop systems. This expands the functional repertoire of the genus beyond its established roles in salt tolerance^10,19–22^ and provides the first proof-of-concept that immune-associated functions from halophytic microbiomes can be transferred into glycophytic crop systems. The ability of *Kushneria* to persist and function under saline conditions further highlights their potential as resilient microbial inoculants in environments where conventional beneficial microbes often lose their biocontrol efficacy^1–4^.

Our multi-omics analyses indicate that *Kushneria*-mediated protection is associated with a distinct immune reprogramming response involving the activation of core NLR receptors, including a ZAR1-like paralogue (Solyc07g053010.3). On the other hand, the ZAR1 orthologue (Solyc02g084890.3) of *Arabidopsis thaliana* ZAR1 (HopZ-Activated Resistance 1) remained unaffected in our transcriptomic analysis. This response was accompanied by transcriptional activation of salicylic acid- and jasmonic acid-associated defense pathways and downregulation of photosynthesis-and translation-related genes, suggesting widespread transcriptional reprogramming during early colonization. Importantly, these immune-associated responses coincided with disease protection under both controlled and field conditions without evident adverse effects on plant performance in the tested conditions. Although NLR activation has been reported in selected beneficial interactions^23,24^, the regulatory architecture underlying such responses remains poorly resolved.

By integrating mRNA, miRNA and lncRNA datasets, we identified a previously uncharacterized long non-coding RNA, Solyc10r048010.1, as a highly connected regulatory hub predicted to interact with multiple induced NLR loci. This lncRNA represents a predicted target of sly-miR167-3p. Treatment with *Kushneria* showed a reduction of the miRNA’s levels, thus alleviating the repression activity and leading to upregulation of the lncRNA and its downstream targets, including the NLRs. Interestingly, among the targets of the novel-sly-miR101-5p, which was upregulated, was one of the five NLRs. This miRNA has high sequence similarity to miR166 from various plant species, and may be a regulator of the NLR homeostasis, as similarly been shown for the miR482/2118 family of miRNAs^24,25^. These findings identify a candidate RNA–NLR regulatory framework associated with *Kushneria*-induced immune reprogramming. Although functional validation will be required to establish causality, this framework provides a tractable mechanistic hypothesis for transferable microbiome-mediated immunity.

The ecological and agricultural relevance of these findings is underscored by the robust field performance of *Kushneria*. Comparative genomics revealed extensive biosynthetic novelty and conserved rhizosphere-competence traits likely associated with persistence in challenging environments. Lipopeptides and other secondary represent candidate determinants of the observed immune priming, as it has been shown for *Bacillus amyloliquefaciens* FZB42 in *Arabidopsis thaliana*^26^. Such traits may confer a physiological advantage over conventional inoculants, whose efficacy frequently declines under abiotic stress. Consistent performance of *Kushneria* across a 77-day field trial, together with protective effects against multiple pathosystems, highlights the translational potential of halophytic endophytes for crop protection under saline agricultural conditions. Together, our findings establish *Kushneria* as a proof-of-concept system demonstrating that halophytic microbiomes can harbour transferable host-protective functions that remain effective following transfer into glycophytic crop systems. Importantly, our findings suggest that the biological relevance of crop microbiome engineering depends not only on natural host association, but also on the transferability of beneficial functions across ecological boundaries. More broadly, this work provides a conceptual framework for exploring extreme-environment microbiomes as reservoirs of transferable functions for crop microbiome engineering and the development of climate-resilient cropping systems.

## Methods

### Samplings, microbial isolation and DNA extraction for WGS

Sampling was made from Chrysi island, located in Crete, Greece in 2021. Samples were collected with sterile tools, bagged separately, and transported at 4 °C to the laboratory for processing. In total, 10 halophytes were collected, from which the endophytic bacteria were isolated from surface-sterilized roots and leaves with a previously described method^14^. *Kushneria* spp. were identified via 16S rRNA gene sequencing and then a DNA extraction followed for whole genome sequencing, based on a publicly available protocol by A. Kiledal and J. A. Maresca^27^ modified from a method by T. Wecke.

### Genome sequencing, assembly and annotation

DNA from all the isolates was sequenced in the BGI Genomics center with both short and long reads, using the sequencing platforms DNBSEQ (sequencing read length PE150) and Cyclone, respectively. Information about the sequencing data is available in Extended Data Table 1. After short reads’ sequencing, the raw reads were filtered. Data filtering included removing adapter sequences, contamination and low-quality reads from raw reads, using the SOAPnuke^28^ developed by BGI with software filter parameters: "-n 0.01 -l 20 -q 0.4 --adaMis 3 --rmdup --minReadLen 150”. Similarly, the long reads were filtered, for removal of adaptors, reads with mean quality lower than 7 and shorter than 1,000, using the software porechop (Version 0.2.4 and default parameters).

Hybrid assemblies were performed using Unicycler (v0.5.0)^29,30^ with default parameters. The structural statistics and functional completeness of the assemblies were evaluated using QUAST v5.0.2^31^ with default options, BUSCO v5.8.0^32^ with the *Pseudomonadota* database (pseudomonadota_odb12) and CheckM2 v1.1.0^33^ with default parameters.

Genome annotations and genome maps were generated using Bakta v1.12.0 with its full database v6.0^34,35^. RAST^36^ was also used to determine if secretion systems were annotated in the sequenced genomes. Plasmid detection was performed with RFPlasmid v0.0.18^37^ using the parameter – species *Pseudomonas*, as *Pseudomonas* represented the closest available genus to *Kushneria* within the Phylum *Pseudomonadota*.

To identify potential type-III secretion system and/or type-III secreted effectors across the assembled genomes, the predicted proteomes were analyzed using the machine learning-based pipeline Effectidor II^38^. Predictions were executed using default parameters, with candidate effectors automatically cross-referenced by the tool against its internal database of curated, experimentally validated T3SEs.

The sequences of the 9 *Kushneria* isolates were uploaded to the European Nucleotide Archive (ENA) under the PRJEB105379 study accession number. The metadata of the samples follow the MIxS-host-associated checklist of the Genomic Standards Consortium^39^.

### Taxonomic and phylogenetic analyses

The genome sequence data were uploaded to the Type (Strain) Genome Server (TYGS), a free bioinformatics platform available under https://tygs.dsmz.de, for a whole genome-based taxonomic analysis^40^. The analysis also made use of recently introduced methodological updates and features^41,42^. Information on nomenclature, synonymy and associated taxonomic literature was provided by TYGS’s sister database, the List of Prokaryotic names with Standing in Nomenclature (LPSN, available at https://lpsn.dsmz.de)^41,42^. The results were provided by the TYGS on 2026-02-17. Because the TYGS database did not contain all NCBI listed sequenced genomes of this genus, we manually uploaded the genomes that were missing. These are namely our 9 sequenced isolates (SRL376, SRL377, SRL383, SRL386, SRL403, SRL411, SRL412, SRL423 and SRL426), as well as *Kushneria* sp. EE4, *Kushneria* sp. TE3, *Kushneria* sp. AK178, *Kushneria avicceniae* CCM 7396, *Kushneria* sp. 4-2069, *Kushneria* sp. Sum13 and *Kushneria aurantia* CCM 7415. The TYGS analysis was subdivided into the following steps: Determination of closely related type strains, pairwise comparison of genome sequences, phylogenetic inference and Type-based species and subspecies clustering.

Determination of closest type strain genomes was done in two complementary ways: First, all user genomes were compared against all type strain genomes available in the TYGS database via the MASH algorithm, a fast approximation of intergenomic relatedness^43^, and, the ten type strains with the smallest MASH distances chosen per user genome. Second, an additional set of ten closely related type strains was determined via the 16S rDNA gene sequences. These were extracted from the user genomes using RNAmmer^44^ and each sequence was subsequently BLASTed^45^ against the 16S rDNA gene sequence of each of the currently 24127 type strains available in the TYGS database. This was used as a proxy to find the best 50 matching type strains (according to the bitscore) for each user genome and to subsequently calculate precise distances using the Genome BLAST Distance Phylogeny approach (GBDP) under the algorithm ’coverage’ and distance formula d5^46^. These distances were finally used to determine the 10 closest type strain genomes for each of the user genomes.

For the phylogenomic inference, all pairwise comparisons among the set of genomes were conducted using GBDP and accurate intergenomic distances inferred under the algorithm ’trimming’ and distance formula d5^46^. 100 distance replicates were calculated each. Digital DDH values and confidence intervals were calculated using the recommended settings of the GGDC 4.0^42,46^. The resulting intergenomic distances were used to infer a balanced minimum evolution tree with branch support via FASTME 2.1.6.1 including SPR postprocessing^47^. Branch support was inferred from 100 pseudobootstrap replicates each. The tree was rooted at the midpoint^48^, visualized with PhyD3^49^ and processed with Inkscape v.1.4.3. The type-based species clustering using a 70% dDDH radius around each of the 29 type strains was done as previously described^40^. Subspecies clustering was done using a 79% dDDH threshold as previously introduced^50^.

### Pangenome analysis

All the newly sequenced *Kushneria* isolates, as well as all sequenced strains deposited in NCBI from the genus (Access date: 05/01/2026, Ext. Data Table 5) were processed for a pangenome analysis using anvi’o v8^51,52^. Genomic and functional annotation was conducted on genomes storage databases generated with anvi-gen-genomes-storage using anvi-run-hmms (with the flags: -I Bacteria_71 and --also-scan-trnas)^53^, anvi-run-ncbi-cogs^54^, anvi-run-kegg-kofams^55^, anvi-run-pfams^56^ and anvi-run-scg-taxonomy^57,58^. The anvi-pan-genome command was used with --minbit 0.5, --mcl-inflation 10, and --use-ncbi-blast^51,52^. Gene cluster presence/absence data were extracted from the SQLite PAN.db files of each pangenome analysis using sqlite3 (version 3.45.3), and post-processed with gawk (GNU Awk 5.3.0) to count gene cluster occurrences per genome. The graphs were created in R (version 4.5.1)^59^ using tidyverse v2.0.0^60^, and UpSetR v1.4.0 [60] and the heatmap using pheatmap v1.0.13^61^. The final figure including the heatmap and the barplot was configured using patchwork v1.3.2^62^.

### Primary and secondary metabolism

Metabolic potential was inferred from KEGG module completeness profiles derived from the pangenome analysis. Module-level outputs from all *Kushneria* spp. genomes were parsed and merged in R (version 4.5.1)^59^ using tidyverse v2.0.0^60^. Genome names were manually curated to ensure consistency across datasets. A binary presence–absence matrix was constructed, whereby a module was considered present if stepwise completeness was ≥ 0.8 or if the module was classified as complete based on pathwise reconstruction. Missing values were treated as absence. Modules conserved across all genomes or absent from all genomes were excluded, retaining only variable modules. Module identifiers were annotated using KEGG module names.

The resulting matrix was visualized as a heatmap using pheatmap v1.0.13^61^, with hierarchical clustering of both genomes and modules based on correlation distance. For the identification of secondary metabolites, only the complete genomes of *Kushneria* genus (Ext. Data Table 5) were screened for biosynthetic gene clusters (BGCs). We did not characterize the genomes that were not complete (contig or scaffold assembly), as the missasembly could affect our results. All assemblies were uploaded to the antiSMASH version 8.0.1 platform for BGC prediction^63^. Similarly, for the identification of rhizosphere-competence-related catabolic gene clusters (rCGCs), all complete genomes were uploaded to the rhizoSMASH^18^ WebApp (https://rhizosmash.bioinformatics.nl/). For both, the detection strictness was set “relaxed”, and all additional features were enabled. Information about the BGC and rCGC groups of each isolate and their similarity confidence with known BGCs was manually collected and organized into categories. Specifically, for BGCs, all NI-siderophores were grouped into the same category, and similarly terpene and terpene-precursor, into the category named “terpene”. BGC and rCGC plots were created using ggplot2 v4.0.2^64^, tidyverse v2.0.0^60^ and patchwork v1.3.2^62^.

### Pathogen inhibition assays

#### Bacterial cultures

For all the *in vitro* and *in planta* assays, the *Kushneria* spp. examined were cultured in Nutrient Agar (NA, 0.5% w/v bacto-peptone, 0.3% w/v yeast extract and 0.5% w/v NaCl, pH 7.4) or Nutrient Broth (NB, for liquid cultures) supplemented with 3% NaCl (w/v), which establishes optimal hypertonic conditions for the growth of halophilic *Kushneria* strains. For the *in-planta* assays, bacterial cells were subsequently harvested by centrifugation at 4,000 × g for 10 min and were washed twice with sterile ultrapure water before use. For other bacteria used, such as *Streptomyces* sp. SRL307, NB or NA was used and in specific for *in planta* assays, the bacterial culture was washed twice before use, as previously described.

#### In vitro assays

The *Kushneria* spp. isolates were screened in confrontation assays together with the phytopathogenic fungi *Fusarium oxysporum* f.sp. *radicis*-*lycopersici* (*Forl*), *Botrytis cinerea* and the oomycete *Phytopthora nicotianae*. The pathogens were freshly grown in potato dextrose agar (PDA) medium (otherwise stated) and plugs were used for the *in vitro* assays. The confrontation tests were made in PDA medium, unless otherwise stated. For *Forl*, *in vitro* dual-culture antagonism assays were executed on two distinct cultural media: (i) Nutrient Agar (NA) supplemented with 3% NaCl (w/v), which establishes optimal hypertonic conditions for the growth of halophilic *Kushneria* strains and (ii) Potato Dextrose Agar (PDA), serving as a standard nutritional growth medium for the fungal pathogen. All plates were incubated in triplicate at 27 °C for 10 days, except in the case of *Phytopthora nicotianae* where 6 replicates were included in the assay to reduce variability.

### In-planta assays

#### Fusarium oxysporum f. sp. radicis-lycopersici infections

The phytopathogenic fungal isolate *Fusarium oxysporum* f. sp. *radicis-lycopersici* (*Forl*) was kindly provided by Dr S.E. Tjamos (Laboratory of Phytopathology, Agricultural University of Athens). The fungal isolate was cryopreserved at - 80 °C by freezing a conidial suspension in 25% v/v aqueous glycerol^65^. Prior to experimental use, the fungus was transferred to potato dextrose agar (Merck, Darmstadt, Germany) and incubated at 27 °C for seven days. Tomato seeds (*Solanum lycopersicum* L., cv. ACE) were surface-disinfected (1.0% v/v NaOCl for 1–2 min), rinsed, and sown in a sterile compost (Potground; Klasmann). At the cotyledon stage, uniform seedlings were transplanted into 9-cm pots and maintained in a greenhouse under controlled environmental conditions (25 °C ± 2 °C, 18/6 h light/dark photoperiod). Bacterial inoculum (1×10^8^ CFU mL^-1^) was applied 15 days post-transplanting at the 2–3 true-leaf stage via root drenching, by administering 20 mL of the adjusted bacterial suspension per pot.

For fungal inoculum, *Forl* was cultured in liquid sucrose sodium nitrate (SSN) medium inoculated with agar plugs (0,5 × 0,5 cm) excised from the margins of the actively growing fungal colonies and incubated at 27 °C under continuous shaking at 120 rpm for three days. The culture was filtered through sterile gauze, centrifuged at 8,000 × g for 10 min and resuspended in sterile distilled water to a final density of 10^6^ conidia mL^-1^. Fungal inoculation was executed three days after the bacterial application via root drenching, by administering 20 mL of the conidial suspension per plant.

Disease symptoms (chlorosis, wilting, stem base rot) were monitored twice weekly for 30 days post-inoculation (dpi). At 30 dpi, plants were harvested to record shoot and root fresh weights, and vascular discoloration was visually assessed via longitudinal stem sections.

To confirm Koch’s postulates and endophytic colonization, stem sections were halved: one half was used for microbial re-isolation on PDA (with antibiotics) and NA (with 3% w/v NaCl and 0.01% w/v cycloheximide), while the other half was flash-frozen in liquid nitrogen and stored at −80 °C.

The area under the disease progress curve (AUDPC) was calculated by the trapezoidal integration method^66^.

#### Botrytis cinerea infections

For the *in planta* and *ex vivo* experiments, *Solanum lycopersicum* Belladona F1 plants and *Solanum lycopersicum* Lobello F1 biological fruit were used, respectively. The plants (all the leaves of the plants) or the fruit were immersed in the bacterial suspension (approx. 10^8^ CFU mL^-1^ of isolates of *Kushneria* spp. or *Streptomyces* SRL307) or sterile distilled water (mock). 3 days post treatment, the plants or the fruit were infected (artificially inoculated) with the fungus *Botrytis cinerea* isolate B.14.9^67^. Specifically, 4 leaves of five plants (in total 20 replicates) or 21 fruit from 3 individual plants (7 fruit/plant) were slightly injured with a sterile needle and infected with fresh culture of the fungus (mycelial plugs of 5 mm diameter obtained from the actively growing margins of fresh *B. cinerea* cultures). The treated plants/fruit were incubated in separate, high-humidity plastic boxes, maintained in a growth chamber at 20 °C and 12 hours photoperiod. Disease severity was evaluated daily by measuring the diameter of the rot (in mm), surrounding the fungal inoculum. The AUDPC was calculated the last day of the experiment (3^rd^ day for the *in planta* and 6^th^ day for the *ex vivo*) according to the trapezoidal integration method^66^.

#### Phytopthora nicotianae infections

Tomato plants (*Solanum lycopersicum*) hybrid Vitara (F1) were used for the *in-planta* assay with the oomycete pathogen *Phytopthora nicotianae*. Seeds were sown individually in plastic pots containing a standardized commercial soil substrate and maintained under controlled greenhouse conditions (16 h light and 8 h dark photoperiod and 65% relative humidity). Fifteen days after sowing, plants received an initial root drench treatment with 10^8^ CFU mL^-1^ of the *Kushneria* suspension. Five tomato plants were treated with the bacterial suspension, and five plants were used as control treated with an equal volume of distilled water. A second application was performed one week later.

When plants reached approximately 30 cm in height, they were inoculated at the stem base with the oomycete *Phytophthora nicotianae*. A small incision of 0.5 cm was performed on the surface of the stem base of the plants using a sterile scalpel and a 6 mm mycelium plug from an actively growing culture of the oomycete was inserted into the wound.. The inoculation area was covered with parafilm to maintain the inoculum in place and to ensure the necessary humidity for infection. In control plants the same incision was performed and covered with parafilm, but without inoculum, in order to maintain identical conditions.

#### Podosphaera xanthii infections

Cucumber (*Cucumis sativus* L.) F1 hybrid Proteas (Syngenta AG, Switzerland) was used. Seeds were sown individually in 15-cm plastic pots containing sterile peat substrate for germination. One week post-emergence, seedlingswere transplanted into 2 L pots. All plants were grown under controlled greenhouse conditions (16-h light/8-h dark, 70% relative humidity, 22 °C). Plants were arranged in a completely randomized, block design with 5 blocks (replicates) comprising a total of 25 plants, including mock (H_2_O), pathogen alone and pathogen along with either SRL376, SRL412 or SRL426 (5 plants/treatment).

Two weeks after sowing, the foliage of the plants was sprayed with different *Kushneria* isolates (approx. 10^8^ CFU mL^-1^). A second application was performed one week later.

Artificial inoculation with *Podosphaera xanthii* conidia was carried out on 22-day-old soil-grown plants (24 hours after the second treatment) by spraying them with a conidial suspension adjusted to 1.50×010^5^ conidia ml^−1^. Disease severity (expressed as the percentage of infected leaf area) was assessed visually according to EPPO STANDARD PP1-057-3 “Powdery mildew of cucurbits and other vegetables” (EPPO, 2005).

*Ex vivo* bioassays were carried out on cucumber leaf disks. Sterile, circular filter papers were placed into 9 cm Petri dishes and moistened with 5 mL of sterile distilled water to maintain humidity. Leaf discs (15 mm diameter) were obtained from the third or fourth leaf. The disks were dipped for a few seconds into each *Kushneria* isolate suspension using sterile forceps, allowed to air-dry on sterile filter paper and subsequently transferred to Petri plates. A 20 μL droplet of the *P. xanthii* conidial suspension (1.50×010^5^ conidia ml^−1^) was applied to the center of each disk. The plates with the disks were transferred to a growth chamber with controlled conditions (16 h light / 8 h dark, 70 % relative humidity, 22 °C). Disease severity in inoculated discs was estimated visually as the percentage of infected leaf surface area.

### Field trial assay

The field assay was performed in the area of Mesaras (Ampelouzos) in South Crete, using hybrid *Solanum lycopersicum* cv. Elpida F1. The plants had been planted in five double-row plots, 49 days prior to the first treatment. For each treatment, 100 plants were used (20 plants per double-row plot), with untreated, non-inoculated buffer plants left between treatment plots to prevent drift and cross-contamination. .

Specifically, for the preparation of the bacterial suspensions, 2 L of concentrated bacterial suspension of *Kushneria* sp. SRL386 (concentration 1.65×10^8^ CFU mL^-1^ at a dilution of 1/10) were diluted in water in a 10 L sprayer, while for the control intervention (Control-) 2 L of liquid nutrient substrate of sterile TSB was diluted. In addition, a commercial preparation (X) was used as a reference product. For the commercial product, 25 gr were diluted in 10 L of water, according to the manufacturer’s instructions. A second spray treatment followed after two weeks.

To monitor the progression of the natural *B. cinerea*-causing disease, measurements were obtained every 2-3 weeks, recording the number of infected leaves, stems and/or fruits from the plants after each treatment. The day 0 was set the day of the second spray treatment and the experiment ended at day 77. Efficacy evaluation was performed according to EPPO Standard PP1/054 (3)

### Statistical analyses and graphs of *in vitro*, *in-planta* and field assays

The statistical analyses of all the *in vitro* and *in planta* experiments, including the field assay, were made with one-way ANOVA, followed by a post-hoc Tukey, in R, using the Base R. For the boxplots of all *in vitro* and *in planta* assays and the barplot of field assay were conducted with ggplot2 (v.3.5.2^64^), ggpubr (v.0.6.1^68^) and rstatix (v.0.7.3^69^). The panels of the figures were composed with patchwork (v.1.3.1^62^).

### Priming treatment, RNA extraction cDNA synthesis and qRT PCR

#### Priming treatment

*Solanum lycopersicum* cv. Micro-Tom plants were grown under controlled greenhouse conditions (16-h light/8-h dark, 70% relative humidity, 24 °C). One-month-old plants were subjected to root drenching treatments with *Kushneria* sp. SRL386, *Streptomyces* sp. SRL307 or sterile ddH_2_O (mock). Specifically, to prepare the inoculum, *Streptomyces* sp. SRL307 was cultured in standard Nutrient Broth (NB), while *Kushneria* sp. SRL386 was grown in NB supplemented with 3% (w/v) NaCl. Bacterial cells were harvested by centrifugation, washed twice, and resuspended to a final concentration of 10^8^ CFU mL^-1^. Plants were root drenched with 15 mL of each bacterium (or sterile ddH_2_O) (3 replicate plants per treatment). Leaf samples were harvested from the treated plants at 0, 24, 48, 72 and 168 h post inoculation.

#### RNA extraction, cDNA synthesis and qRT-PCR

Tomato leaves of each treatment were cut and immediately frozen in liquid nitrogen. Leaf tissue was grounded to a fine powder, with liquid nitrogen using a sterile mortar and pestle. 100 mg of leaf tissue was used to extract total RNA, with TRIzol reagent.The extracted RNA was purified via phenol-chloroform extraction and precipitated using a 1:1 ratio of isopropanol and high-salt solution (0.8 M sodium citrate and 1.2 M NaCl) . RNA concentration was measured in NanoDrop™ Spectrophotometer (Thermo Fisher Scientific Inc., Waltham, Massachusetts, United States) and visualized its integrity in a non-denaturing agarose gel.

To remove any residual genomic DNA, 10 ug of total RNA was treated with DNAseI (Roche, 04716728001), for 1 hour at 37 °C. The cDNA synthesis was performed with the use of engineered M-MLV reverse transcriptase basic kit (EnzyQuest, RN012S) and 1 ug of total RNA, according to the manufacturer’s instructions. No-RT samples were also prepared (without the addition of M-MLV RT), to ensure that no DNA had remained after DNAseI treatment.

qRT-PCR analysis was performed with the use of KAPA SYBR^®^ Fast Master Mix (2x) Universal kit (KK4602) and with a dilution of 1/50 of cDNA samples (corresponding to 1 ng) and according to the manufacturer’s instructions. The primers with their sequences used are listed in the Extended Data Table 13.

### Transcriptome sequencing and analysis

#### Transcriptome sequencing

RNA samples were sent to BGI Genomics for sequencing of mRNA, long non-coding (lncRNAs) and small RNAs (sRNAs). For all of them, the DNBSEQ platform was used with PE150, PE100 and PE50 for mRNAs, lncRNAs and miRNAs, respectively.

#### Transcriptome analysis

The transcriptome analysis of all RNAs sequenced was perfomed by BGI. For mRNAs, Bowtie2^70^ (v.2.2.5, parameters -q --sensitive --dpad 0 --gbar 99999999 --mp 1,1 --np 1 --score-min L,0,-0.1 -p 16 -k 200) was used to map the clean reads to the reference trascriptome (Solanum_lycopersicum_solgenomics.net_ITAG4.0). Then RSEM^71^ (v.1.2.8, parameters-p 8 --forward-prob 0 --paired-end) was used to calculate the gene expression level of each sample. For sRNAs, Bowtie2^70^ (parameters: -q -L 16 --phred64 -p 6) was used to align clean reads to the reference set and other small RNA databases, except that Rfam was aligned using cmsearch (parameters: --cpu 6 –noali). For lncRNAs, Bowtie2^70^ (parameters: -q --sensitive --dpad 0 --gbar 99999999 --mp 1,1 --np 1 --score-min L,0,-0.1 -I 1 -X 1000 --no-mixed --no-discordant -p 1 -k 200) was used to map clean reads to the reference sequence, and then used RSEM^71^ (v. 1.2.12, parameters: --forward-prob 0) to calculate the expression levels of transcripts.

For the differential expressed genes (DEGs) of all RNAs sequenced, Dr. Tom platform was used with settings of |log2FC| >= 1 and Qvalue <= 0.05 (for sRNAs the Qvalue was set to 0.001). The detection was performed with DEseq2 using the method described in Michael I, et al.^72^ for mRNAs and lncRNAs, and DEGseq^73^ for sRNAs.

To allocate the DEGs to different metabolic pathways, the up- and down-regulated transcripts were divided and selected to perform enrichment in KEGG database. For the GSEA analysis in the KEGG database, the unit of measure was the normalized read count and the filtering threshold was set to 500 max and 15 min size. These analyses, as well as the volcano plots were generated in Dr. Tom platform of BGI. The plot of GSEA and the volcano plot for lncRNAs were generated in R using ggplot2^64^.

### Prediction of targets of miRNAs and lncRNAs

For the prediction of miRNAs targets, psRNATarget^74^ (Schema V2, 2017 updated) was used via its platform (https://www.zhaolab.org/psRNATarget/home). For this reason, we submitted the differentially expressed miRNAs, as well as the differentially expressed mRNAs, with default settings except of: expectation 4.0, Max. UPE: 25, 1 mismatch allowed in seed region. For the predicted mRNA targets of miRNAs, a plot was generated in R using ggplot2^64^, and colors were given for the KEGG pathway level 2 annotation of every target (where available). Multiple KEGG pathways are shown with division of each bar, with more colors per bar. The targets that are multiply targeted by different miRNAs are shown with a star. For the best visualization of the plot, the total number of mRNA targets is shown below each miRNA.

Additionally, a prediction of the lncRNAs putative targets of miRNAs was performed in psRNATarget^74^, by providing the differentially expressed miRNAs and the differentially expressed lncRNAs, with default settings.

For the prediction of cis targets of lncRNAs, we investigated in a 100 kb proximity of the differentially expressed lncRNAs, which differentially expressed mRNAs are located. We ended up with 26 pairs of DE lncRNAs – DE mRNAs, which we propose as the putative cis targets. For the prediction of trans targets of lncRNAs, we made a Pearson correlation between the DE lncRNAs and DE mRNAs, based on their TPM (transcripts per million). To exclude false positives, we filtered the results with abs. value of r (|r|) > 0.98 and p-value < 0.05. Additionally, we filtered out the cis targets and we ended up with 679 pairs of DE lncRNAs and DE mRNAs. Next, we predicted for all pairs their RNA-RNA interaction and MFE (minimum free energy) and total energy of their interactions, as well as their normalized total energy (ndG = total energy / length of interaction), using RNAplex^75^ from ViennaRNA package^76^. All 679 pairs were kept when we filtered with total energy < -35 kcal/mol & ndG < -0.1.

### Network analysis

For the network analysis, we introduced the DE lncRNAs with their targets (cis and trans) and the DE miRNAs with their targets. For the predicted targets of miRNAs and lncRNAs we filtered with Expectation <= 4.0. The network was constructed in Cytoscape v.3.10.4^77^, with the addition of KEGG pathway Level 2 annotation for all the predicted targets of each lncRNA, in chart pies, separately for up- and down-regulated targets. If a target had multiple KEGG, these were all included in the chart pies. If no KEGG annotation was available, the targets were excluded from the chart pies. The legend for the KEGG annotation was created in R, and merged with the network, using Inkscape v.1.4.3, retrieved from https://inkscape.org.

## Acknowledgments

VAM, TM, SK, MFT, EM and PFS were funded by the European Union - Next Generation EU, Greece 2.0 National Recovery and Resilience plan (TAEDR-0535675). NPA was supported by a PhD scholarship of the Research Funding Program of the Research Committee of the University of Crete managed through the Financial & Administrative Support Unit of the Special Account for Research Funds (ELKE) of the University of Crete (No. 11643), as well as by the European Union - Next Generation EU, Greece 2.0 National Recovery and Resilience plan (TAEDR-0535675). SKS and EAM were funded by both the European Union - Next Generation EU, Greece 2.0 National Recovery and Resilience plan (TAEDR-0535675) and RESEARCH-CREATE-INNOVATE (Grant code/Acronym: Τ2ΕΔΚ-01859/BIOCONTROL), co-financed by the European Union and Greek national funds through the Operational Program Competitiveness, Entrepreneurship and Innovation.

## Author information

Authors and Affiliations

**Department of Biology, University of Crete, 714 09 Heraklion, Crete, Greece**

Vassiliki A. Michalopoulou, Savvas Paragkamian, Nikolaos P. Arapitsas & Panagiotis F. Sarris

**Institute of Molecular Biology and Biotechnology, Foundation for Research and Technology-Hellas, 714 09 Heraklion, Crete, Greece**

Vassiliki A. Michalopoulou, Nikolaos P. Arapitsas & Panagiotis F. Sarris

**Department of Biosciences, College of Life and Environmental Sciences, University of Exeter, Exeter, UK**

Panagiotis F. Sarris

**Department of Agriculture, School of Agricultural Sciences, Hellenic Mediterranean University, Heraklion, Stavromenos, 71410, Greece**

Stefanos K. Soultatos & Emmanouil A. Markakis

**Benaki Phytopathological Institute, Scientific Directorate of Phytopathology, Laboratory of Mycology, 145 61 Kifissia, Attica, Greece**

Theoni Margaritopoulou, Sofia Kosenaki, Eleni Kalogeropoulou & Emilia Markellou

## Authors’ Contribution

VAM and PFS conceived and designed the study. VAM, SP and NPA developed the code. VAM performed the analyses. SKS and EAM performed the *in-planta* assays with *Botrytis cinerea*, as well as the field assay. SK and EK performed the *in-vitro* and *in-planta* assays with Forl. TM, MFT and EM performed the *in-vitro* and *in-planta* assays with *Phytopthora nicotianae* and the *in-planta* assays with *Podosphaera xanthii*. Resources were provided by EM, EAM, EK and PFS. Project’s administration PFS, VAM. Students and Post-Docs supervision by PFS, EM, EK and EAM. Data analysis and results validation by PFS, VAM, TM, EM, EK, EAM. Funding acquisition by PFS. All authors contributed to writing the manuscript and approved the final version.

## Ethics declaration

The authors declare no competing interests.

## Data Availability

The whole genome sequences of the *Kushneria* isolates and the RNA seq data are all deposited in European Nucleotide Archive (ENA) under the accession numbers PRJEB105379 and PRJEB116070, respectively.

## Code Availability

The code and associated documentation are available via GitHub at https://github.com/vilimix/Study-of-Kushneria-spp.

