## Extended Data Figures for "Halophytic endophytes provide transferable immune functions for crop resilience"

### Extended Data Figures

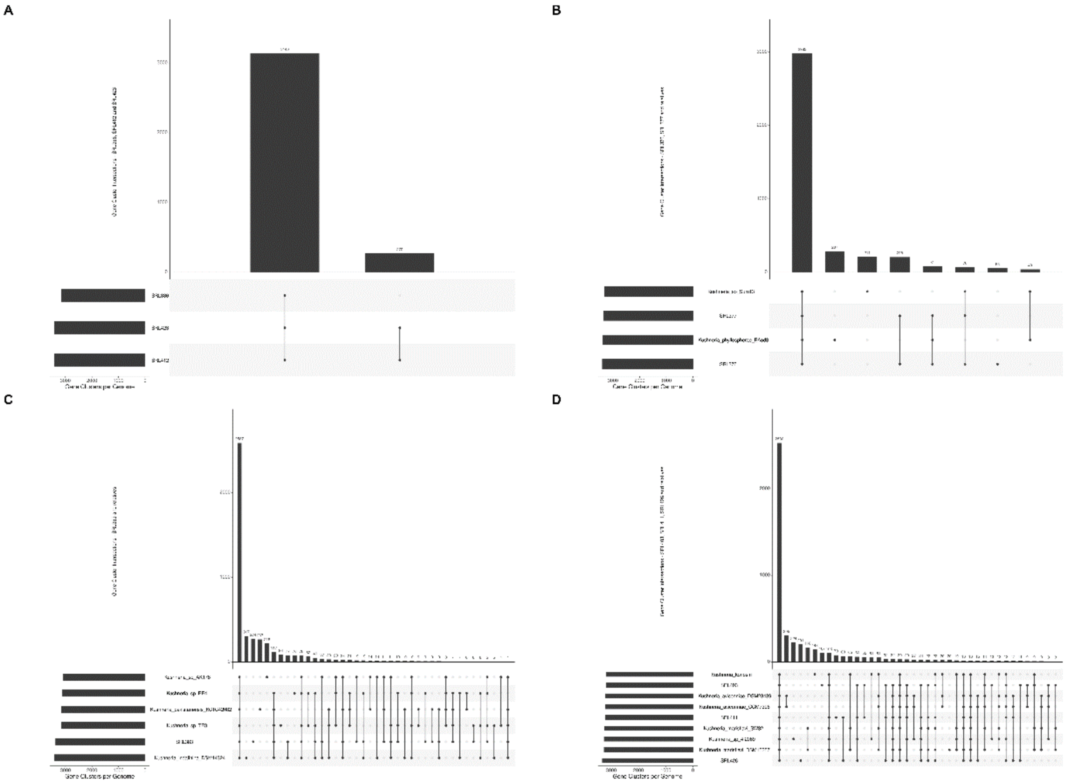


**Extended Data Figure 1. UpSet plots of the different clusters of the *Kushneria* spp. pangenome.**

UpSet plots of the sub-pangenome analyses of the clusters of *Kushneria* spp. **A.** UpSet plot of the cluster of SRL386, SRL412 and SRL423. **B.** UpSet plot of the cluster of SRL376, SRL377 and relatives. **C.** UpSet plot of the cluster of SRL383 and relatives. **D.** UpSet plot of the cluster of SRL403, SRL411, SRL426 and relatives. Horizontal bars (left) represent the total gene clusters per genome, and vertical bars (top) display the number of shared clusters for each intersection (genome combination) indicated by the connected dots below.

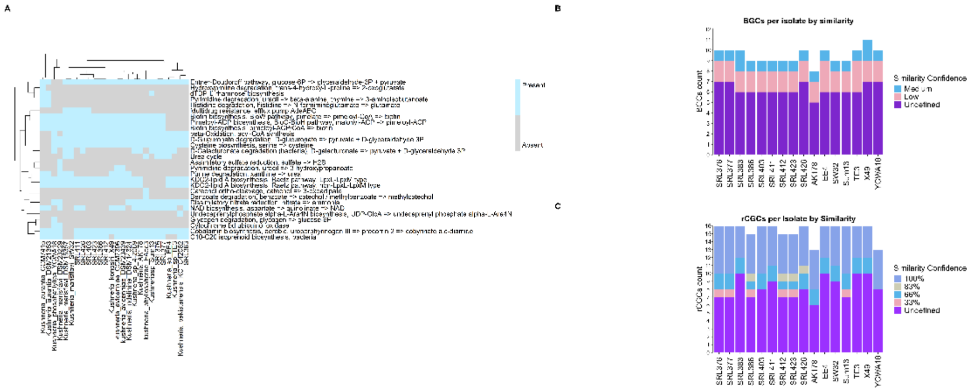


**Extended Data Figure 2. Primary and secondary metabolism distribution of *Kushneria* spp. genomes.**

**A.** Primary metabolism differences of the sequenced genomes of *Kushneria* spp., as calculated by their stepwise and pathwise module completeness. With blue color the pathway shown on the y axis is present, whereas with grey color is absent. **B, C.** Distribution of BGC (**B**) and rCGC (**C**) counts per isolate by similarity confidence. Each stacked bar plot shows the number of BGCs identified in each isolate. Colors indicate the similarity confidence of each BGC/rCGC to known clusters based on the antiSMASH database: Medium, Low, and Undefined (no close matches) for BGCs and 100%, 83%, 66%, 33% and Undefined (no close matches) for rCGCs.

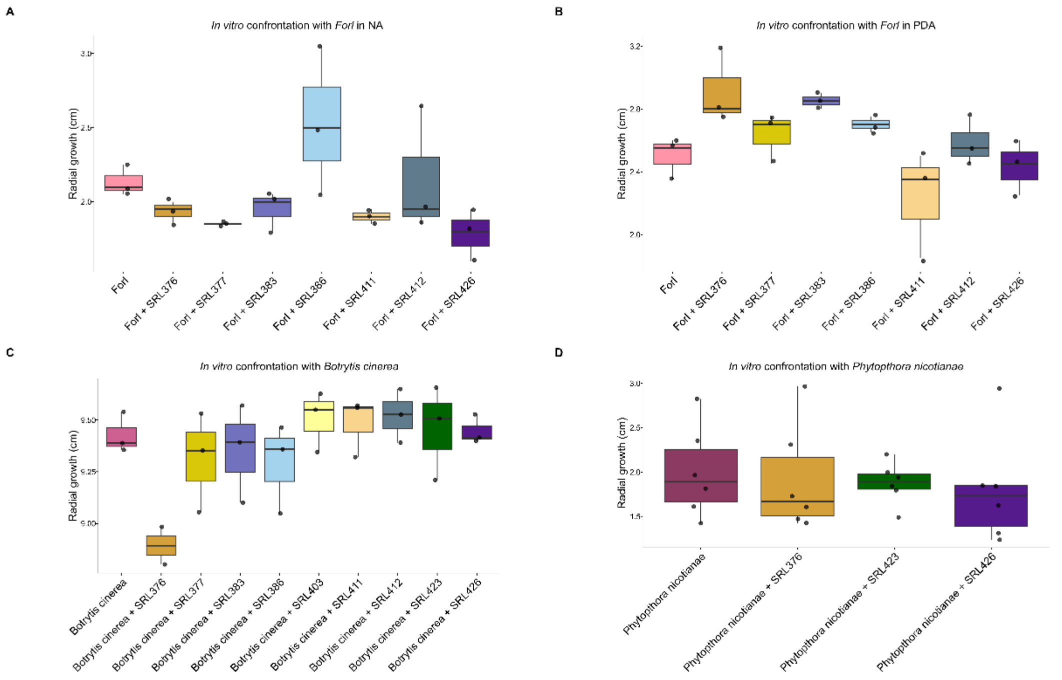


**Extended Data Figure 3. *In vitro* assays of *Kushneria* spp. and different phytopathogens.**

Boxplots showing the radial growth (in cm) of phytopathogens when assaying *in vitro* in the presence of *Kushneria* spp. **A, B.** *In vitro* confrontation of *Fusarium oxysporum* f. sp. *radicis*-*lycopersici* (*Forl*) and SRL376, SRL377, SRL383, SRL386, SRL411, SRL412 and SRL426 in Nutrient Agar (NA) medium (**A**) or PDA medium (**B**). There is no statistically significant reduction of the growth of *Forl* by any *Kushneria* sp., compared to *Forl* only, to any medium tested. **C.** *In vitro* confrontation of *Botrytis cinerea* (*Bc*) and all 9 *Kushneria* spp. sequenced in this study (in PDA medium). There is no statistically significant reduction of the growth of *Bc* in any treatment, compared only to the control. **D.** *In vitro* confrontation of the oomycete *Phytopthora nicotianae* (*Phn*) and SRL376, SRL423 and SRL426 (in PDA medium). There is no statistically significant reduction of the growth of *Phn* in any treatment, compared to the control.

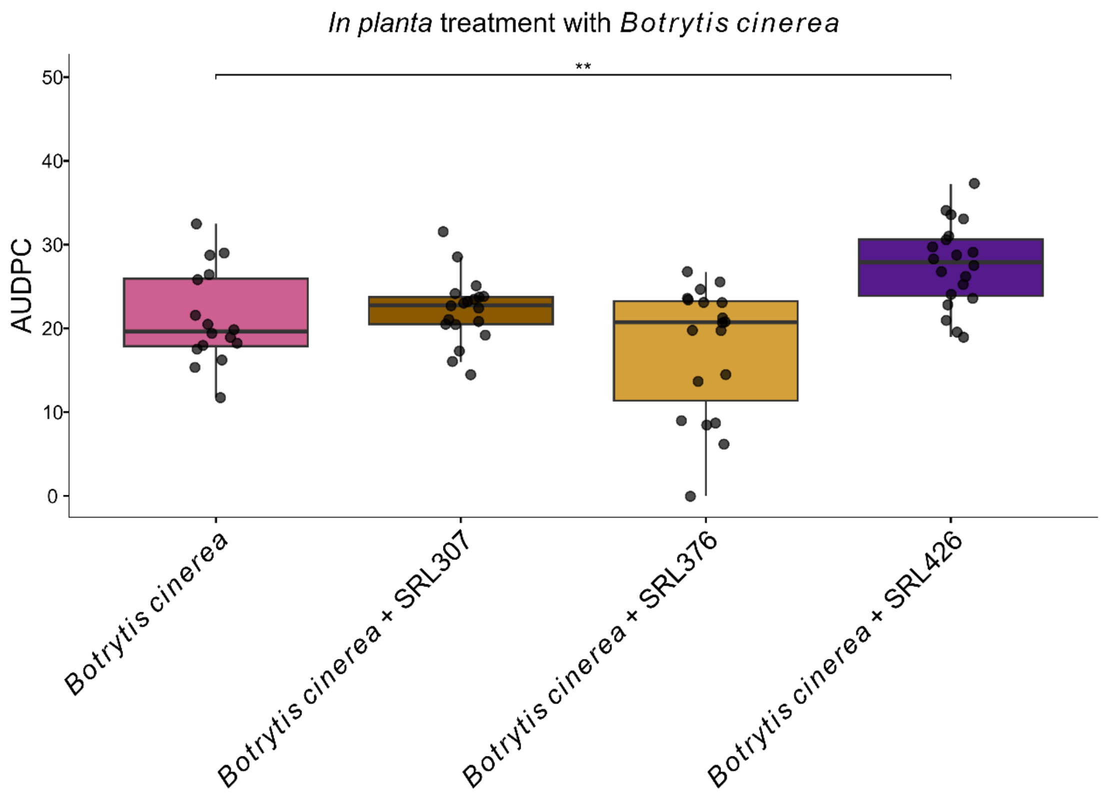


**Extended Data Figure 4. *In planta* assay between *Kushneria* spp. and the *Streptomyces* sp. SRL307 and the phytopathogen *Botrytis cinerea* in *Solanum lycopersicum*.**

Box plot showing the AUDPC caused by *Botrytis cinerea* infection of tomato plants, in the presence of *Streptomyces* sp. SRL307 or the *Kushneria* isolates SRL376 or SRL426. No statistically significant reduction of the AUDPC was observed by any isolate compared to the pathogen alone. The statistical test was Anova and Tukey as a post-hoc with n= 5 plants/treatment and 2-4 infection plugs/plant.

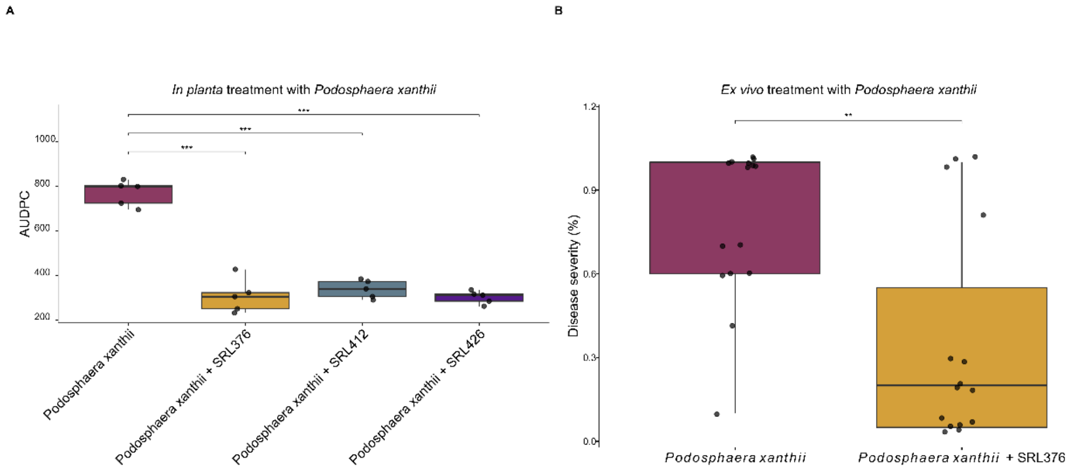


**Extended Data Figure 5. *In planta* assay between *Kushneria* spp. and the phytopathogen powdery mildew in *Cucumis sativus*.**

**A, B.** *In planta* assay of *Kushneria* spp. with *Podosphaera xanthii* in *Cucumis sativus*. **A.** Boxplot showing the AUDPC caused by the fungus *Podosphaera xanthii* along with the treatments of SRL376, SRL412 and SRL426 in cucumber plants. After 27 days of the monitoring of disease severity, the AUDPC was measured, which was decreased in all three treatments, compared to the phytopathogen alone treatment. The statistical test was Anova and Tukey as a post-hoc with n=5 plants/treatment. **B.** Boxplot showing the disease severity (%) caused by the *Podosphaera xanthii* alone or the pathogen and SRL376 in cucumber leaf disks. After 21 days, the disease severity was reduced in the presence of SRL376 treatment. The statistical test was Anova and Tukey as a post-hoc with n= 15 leaf disks/treatment (3 individual plants with 5 leaf disks each).

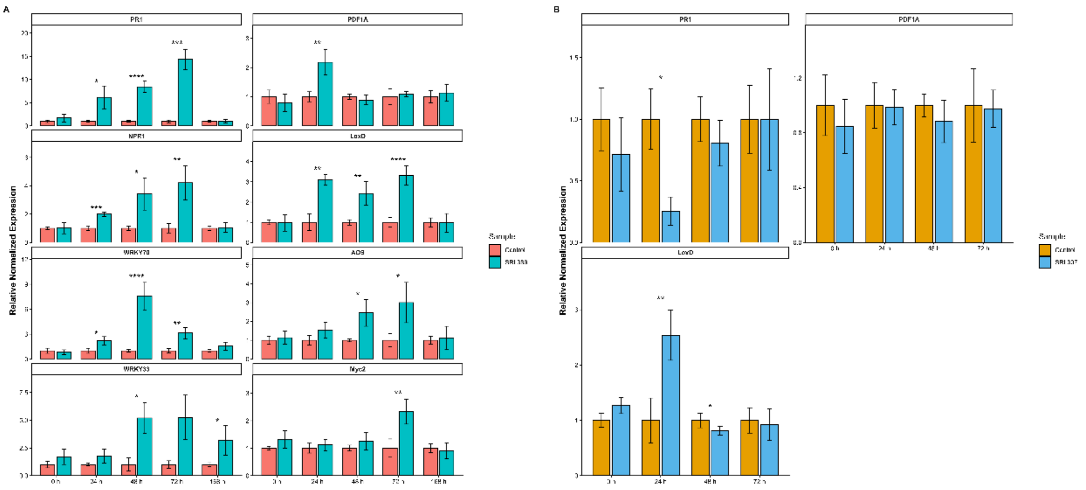


**Extended Data Figure 6. Expression changes of defense genes in tomato plants treated with *Kushneria* sp. SRL386 and *Streptomyces* sp. SRL307.**

Expression changes of the defense genes implicated in salicylic acid signaling and jasmonic acid biosynthesis and signaling: *PR1*, *NPR1*, *WRKY70*, *WRKY33*, *PDF1A*, *LoxD*, *AOS* and *Myc2*. For the qRT-PCRs, RNA from tomato leaves treated with *Kushneria* sp. SRL386 or *Streptomyces* sp. SRL307 was processed, at time points 0, 24, 48, 72 and 168 hours (h) after SRL386 inoculation and from 0 to 72 h in the case of SRL307. **A.** PR1 and NPR1 levels were increased from the 24 until 72 h. WRKY70 was also increased at the same time points, whereas WRKY33 increased later at 48 hours and its levels remained elevated until 168 h. LoxD was increased from 24 until 72 h, while the downstream AOS later at 48 h and its levels remained elevated until 72 h. Myc2 levels were increased at 72 h, while PDF1A only at 24 h and then returned to the control levels. **B.** Regarding the SRL307 treatment, only the PR1 was decreased at 24 h with a simultaneous increase of LoxD levels, while PDF1A expression remained unaffected. Statistical significance stars: *: p-value <0.05, **: p-value <0.01, ***: p-value <0.001 and ****: p-value <0.0001.

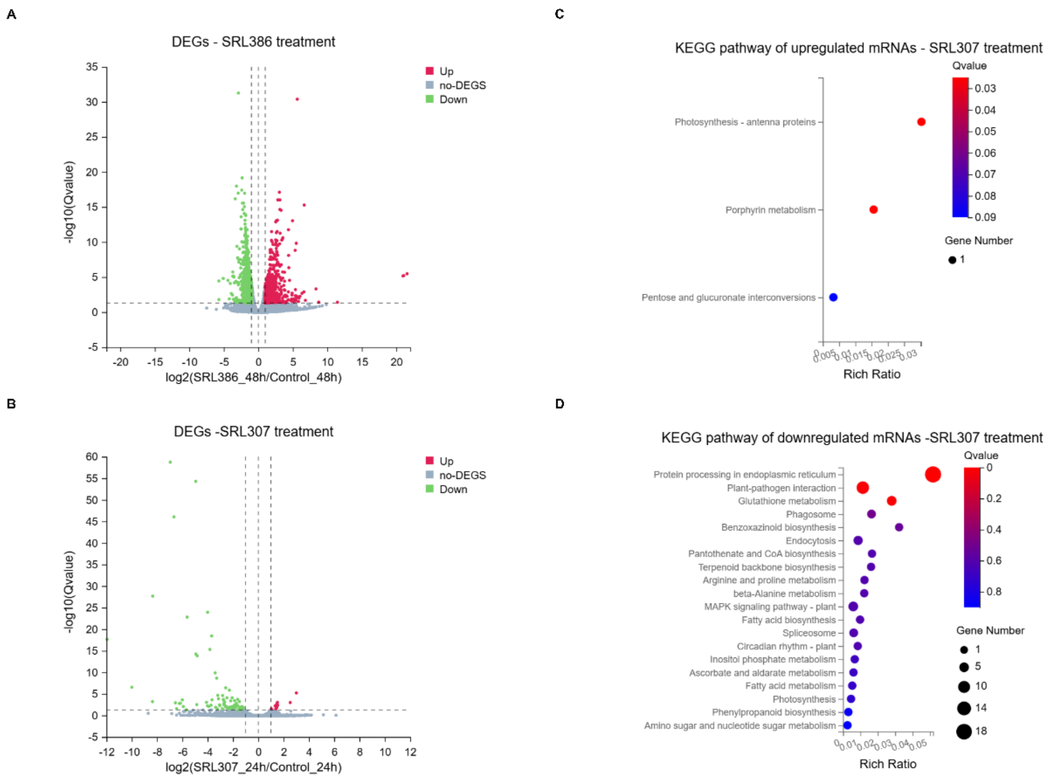


**Extended Data Figure 7. Transcriptomic profile changes of tomato plants treated with *Kushneria* sp. SRL386 or *Streptomyces* sp. SRL307.**

**A, B.** Volcano map showing the differentially expressed mRNAs of the treatment of tomato leaves with *Kushneria* sp. SRL386 at 48 hours **(A)** or *Streptomyces* sp. SRL307 at 24 hours **(B)**, each compared to the relative control treatment ($H_2O$). The absolute value of logFC was set at 1 and Q-value (p-adjusted value) at 0.05. With red color are shown the upregulated genes, with green color the downregulated genes, whereas the grey color represent the non-differentially expressed (no-DEGs) genes. In x and y axes, the log2 and the "-log10" of Q-value are plotted, respectively. The figures are generated in Dr. Tom platform by BGI. **C, D.** An enrichment analysis was made separately for the upregulated (**C**) and downregulated (**D**) genes, according to the KEGG pathway for the SRL307 treatment. A Q-value (correction of P-value) <= 0.05 is regarded as a significant enrichment and is shown by the color gradient. Additionally, the number of genes of each pathway are represented by the size of the bubbles. Most of the upregulated genes were involved in photosynthesis, while the downregulated were implicated in protein processing in ER, plant-pathogen interaction and glutathione metabolism. For more information regarding the DE mRNAs, see Extended Data Table 10.

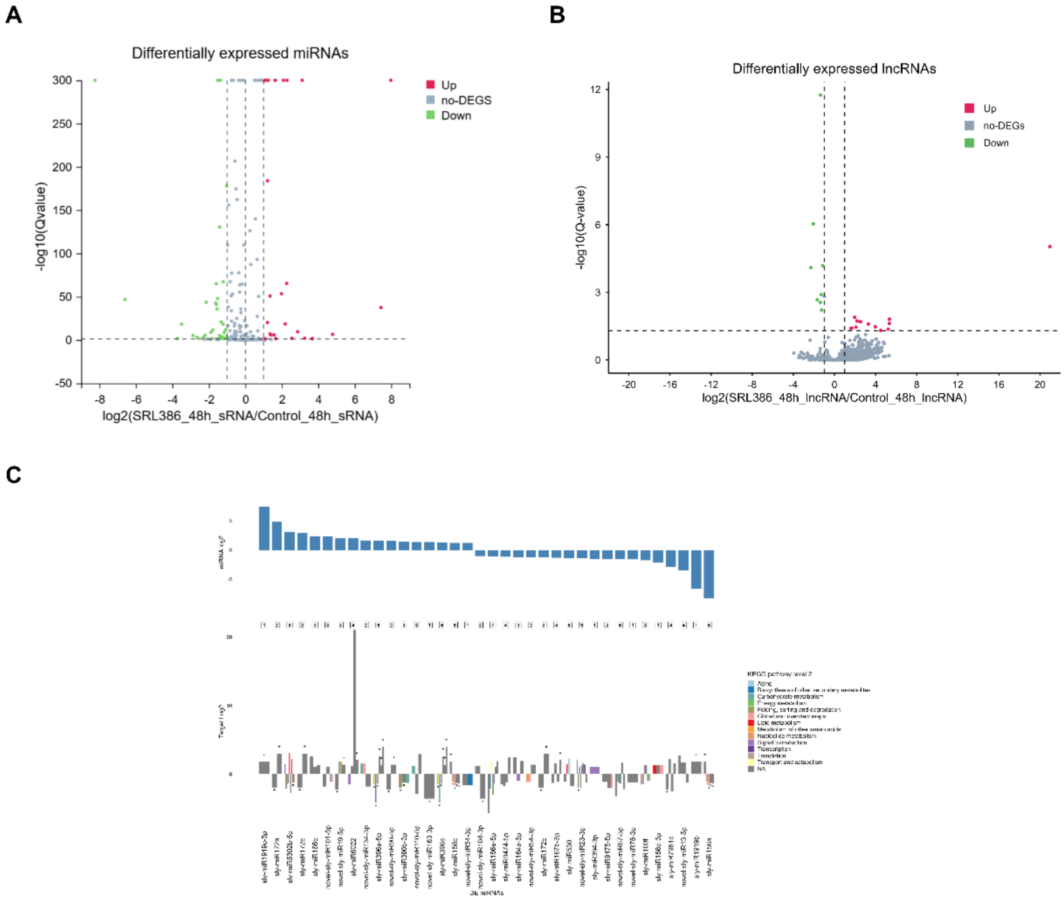


**Extended Data Figure 8. miRNA and lncRNA transcriptomics changes in tomato plants treated with *Kushneria* sp. SRL386.**

**A, B.** Volcano map showing the differentially expressed miRNAs (**A**) and lncRNAs (**B**) of the treatment of tomato leaves with *Kushneria* sp. SRL386, at 48 hours, compared to the control treatment ($H_2O$). The absolute value of logFC was set at 1 and Q-value (p-adjusted value) at 0.05. With red color are shown the upregulated genes, with green color the downregulated genes, whereas the grey color represents the non-differentially expressed (no-DEGs) genes. In x and y axes, the log2 and the "-log10" of Q-value are plotted, respectively. The figure A is generated in Dr. Tom platform by BGI. **C.** A plot with the DE miRNAs (blue color) and their predicted DE mRNAs using the psRNATarget. In total 36 DE miRNAs were predicted to target 98 DE mRNAs. The KEGG pathway term level 2 of each target is shown, where available (grey color represents the NA). Multiple KEGG pathways are shown with division of each bar, with more colors per bar. The total number of targets is shown below each DE miRNA. The stars (*) represent multiply targeted mRNAs. For more information, regarding the annotation of the predicted targets, see Extended Data Table 12.
